# Neurovascular-modulation

**DOI:** 10.1101/2020.07.21.214494

**Authors:** Niranjan Khadka, Marom Bikson

## Abstract

Neurovascular-modulation is based on two principles that derive directly from brain vascular ultra-structure, namely an exceptionally dense capillary bed (BBB length density: 972 mm/mm^3^) and a blood-brain-barrier (BBB) resistivity (*ρ* ~ 1×10^5^ Ω.m) much higher than brain parenchyma/interstitial space (*ρ* ~ 4 Ω.m) or blood (*ρ* ~ 1 Ω.m). Principle 1: Electrical current crosses between the brain parenchyma (interstitial space) and vasculature, producing BBB electric fields (E_BBB_) that are > 400x of the average parenchyma electric field (Ē_BRAIN_), which in turn modulates transport across the BBB. Specifically, for a BBB space constant (λ_BBB_) and wall thickness (d_th-BBB_): analytical solution for maximum BBB electric field (E^A^_BBB_) is given as: (Ē_BRAIN_ × λ_BBB_) / d_th-BBB_. Direct vascular stimulation suggests novel therapeutic strategies such as boosting metabolic capacity or interstitial fluid clearance. Boosting metabolic capacity impacts all forms of neuromodulation, including those applying intensive stimulation or driving neuroplasticity. Boosting interstitial fluid clearance has broad implications as a treatment for neurodegenerative disease including Alzheimer’s disease. Principle 2: Electrical current in the brain parenchyma is distorted around brain vasculature, amplifying neuronal polarization. Specifically, vascular ultra-structure produces ~50% modulation of the average parenchyma electric field (Ē_BRAIN_) over the ~40 μm inter-capillary distance. The divergence of E_BRAIN_ (activating function) is thus ~100 kV/m^2^ per unit average parenchyma electric field (Ē_BRAIN_). This impacts all forms of neuromodulation, including Deep Brain Stimulation (DBS), Spinal Cord Stimulation (SCS), Transcranial Magnetic Stimulation (TMS), Electroconvulsive Therapy (ECT), and transcranial electrical stimulation (tES) techniques such a transcranial Direct Current Stimulation (tDCS). Specifically, whereas spatial profile of E_BRAIN_ along neurons is traditionally assumed to depend on macroscopic anatomy, it instead depends on local vascular ultra-structure.

## Introduction

Vascular responses are ubiquitous across neuromodulation [1–6], but are considered epiphenomena to neuronal stimulation. Common functional imaging techniques measure hemodynamic response (e.g. Arterial Spin Labeling fMRI, H_2_0^15^ PET, SPECT, BOLD fMRI, fNIRS) are interpreted as indexing neuronal activation through neurovascular coupling (NVC). Neurovascular coupling is the mechanism by which increased neuronal activity regulates cerebral blood flow (CBF) to assure that the blood supply of the brain is commensurate to local cellular metabolism [7,8]. The mechanisms of neurovascular coupling are studied to: enhance interpretation of hemodynamic-based imaging techniques [9]; and understand the role of cerebral blood flow and in disease such as hypertension, Alzheimer disease, and stroke [7]. Neurovascular coupling is activated in animals using mechanosensory stimulation [9–11], visual stimulation [12–14], and electrical stimulation of peripheral [15,16] or central axons *distal* to the brain region of interest [17–19]. Stimulation applied directly to a brain region is a special case where brain vasculature can be directly activated [20–22] which: 1) reverses the typical recruitment order of neurovascular coupling, suggesting functional imaging in fact shows direct hemodynamic activation; and 2) resulting in peculiar (supra-physiological) neurovascular changes that suggest novel therapeutic strategies (e.g. metabolic capacity, interstitial clearance).

The brain capillary bed is a dense network of interconnected vessels formed by specialized endothelial cells. The blood-brain-barrier (BBB) is the interface between the blood and brain interstitial fluid. Endothelial cells are sealed together by tight junctions, resulting in an exceptionally resistive BBB. Capillary diameter in the brain is ~10 μm and the average intercapillary distance is ~40 μm [23,24], such that neuronal processes are < 20 μm from the nearest capillary [25]. Moreover, brain capillaries are encased in extracellular matrix proteins and surrounded by specialized neuronal processes and the perivascular end feet of astrocytic glia [26].

Here we consider two consequences of BBB ultra-structure in neuromodulation. First, to what extent does the BBB polarize as a consequence of current crossing between interstitial space and the blood (Principle 1)? Neurovascular coupling and interstitial fluid clearance governs brain health and can be compromised in disease [7]. For example Alzheimer’s Disease (AD) is associated with build-up of misfolded proteins [27,28] and impaired clearance systems [29]. Generally, interstitial fluid clearance is compromised with age [30] which may be linked to the role of clearance during sleep [31]. Interventions enhancing clearance in the brain may treat diverse neurological disorders including of aging [28]. By predicting BBB polarization, Principle 2 provides a substrate for developing neurovascular modulation targeting brain clearance. For example, we proposed tDCS boosts interstitial fluid transport based on BBB electro-osmosis [21].

Second, current flow through the interstitial space is considered insensitive to cellular ultra-structure [32], which has importance consequences for predicting which neuronal elements are stimulated [33]. But, the role of capillaries in distorting current flow is addressed for the first time here (Principle 2). We specifically advance the theory that if microscopic electric field gradients (Activating Function) around neurons created by BBB ultra-structure is larger than that produced by macroscopic tissues changes [34–37], then neuronal stimulation is in fact predicted by the average local electric field [38,39] as convoluted by regional BBB properties. The consequences of this analysis span all forms of brain stimulation including Deep Brain Stimulation (DBS), Spinal Cord Stimulation (SCS), Transcranial Magnetic Stimulation (TMS), Electroconvulsive Therapy (ECT), and transcranial electrical stimulation techniques (tES) such a transcranial Direct Current Stimulation (tDCS).

## Methods

The anatomy of brain vasculature is intractably complex across scales, and current crossing the BBB can exits at neighboring locations or traverse broadly across vascular system, such that macroscopic anatomy may impact microscopic current flow. We overcome this by designing models (e.g. capillary orientation and capillary border boundary conditions) such that assessed variables (e.g. question being asked) were independent of exterior volume dimensions or capillary length. For electric field amplification at the BBB, the models address question regarding the maximum current density crossing the BBB for a given capillary morphology. We also adapt neuron cable theory [40–44] to develop an analytical solution for maximum BBB polarization sensitivity. For addressing neuron polarization amplification by vascular ultra-structure, parallel vessels (with no tortuosity, and region-specific inter-capillary distance) are a conservative model.

### Model Construction and Solution Method

We developed a computer-aided design (CAD) model of BBB ultra-structure to first assess electric field amplification at the BBB (Principle 1) and neuron polarization amplification by vasculature (Principle 2). Different prototypical BBB morphologies were modelled as CAD files in SolidWorks (Dassault Systemes Corp., MA, USA) and imported into Simpleware (Synopsys Inc., CA, USA) to generate an adaptive tetrahedral mesh using a built-in voxel-based meshing algorithm. Mesh density was refined until additional model refinement produced less than 1 % difference in extracellular voltage at the BBB. The resulting model consisted of > 28 million, > 68 million, and > 41 tetrahedral elements for the three exemplary prototypical capillary morphologies: (morphology 1) Semi-circular loop (fixed curvature width) with semi-infinite orthogonal straight segments (Fig. 1A1); (morphology 2) Semi-circular loop (varied curvatures) with semi-infinite parallel straight segments (Fig. 1B1); (morphology 3) Semi-infinite straight tube with variant terminal conditions (Fig. 1C1), and > 38 million, > 29 million, > 45 million, > 68 million, and > 70 million for cortical (Fig. 2A1), white-matter (Fig. 2A2), subcortical (Fig. 2A3), lumbar white-matter (Fig. 2A4), and lumbar grey-matter (Fig. 2A5) vasculature models, respectively.

**Figure 1:**
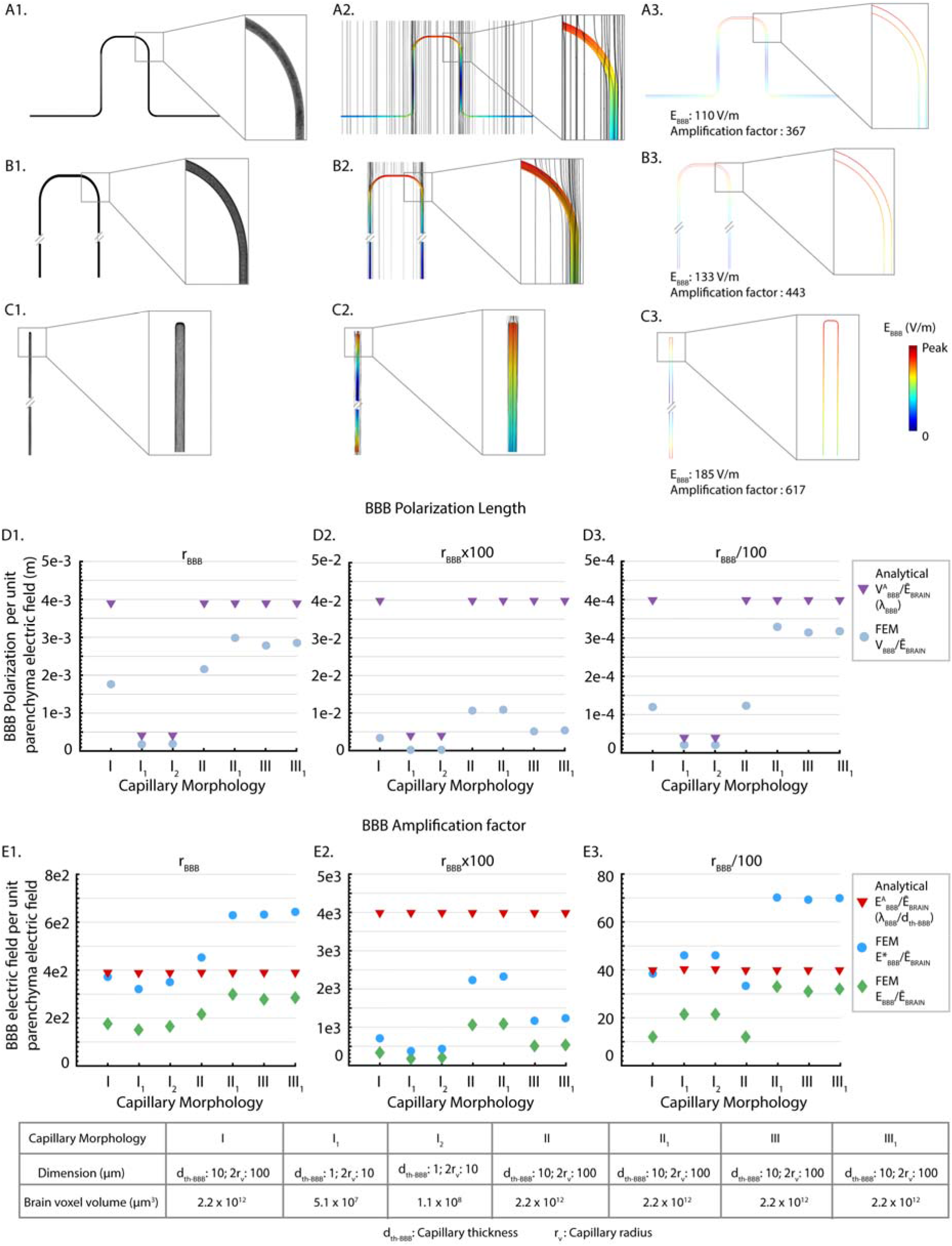
Maximal BBB polarization and electric field amplification across prototypical capillary morphologies compared to analytical maxima. Architecture of three exemplary capillary morphologies (A1) Capillary morphology 1: semi-circular loop (fixed curvature width) with semi-infinite orthogonal straight segments, (B1) Capillary morphology 2: semi-circular loop (varied curvatures) with semi-infinite parallel straight segments, and (C1) Capillary morphology 3: semi-infinite straight tube with tapered end. d_th-BBB_ and 2r_v_ refers to capillary wall thickness and capillary lumen diameter, respectively. Current flow and specifically maximal electric field intensity across the BBB (E_BBB_) was predicted. Capillary morphology 1 include three morphological variations (I, I_1_, and I_2_) with fixed curvature width, but varied d_th-BBB_ (I: 10 μm; I_1_: 1 μm; I_2_: 1 μm) and 2r_v_ (I: 100 μm; I_1_: 10 μm; I_2_: 10 μm). Capillary morphology 2 includes two morphological variations (II and II_1_) with similar d_th-BBB_ (10 μm), 2r_v_ (100 μm) but varied curvature width (II: 1000 μm; II_1_: 200 μm). Capillary morphology 3 includes two morphological variations (III and III_1_) with similar d_th-BBB_ (10 μm), 2r_v_ (100 μm), but varied terminal conditions (III: one end open; III_1_: both ends sealed). Predicted brain current flow pattern (black flux lines) and BBB electric field (false color) are showed for capillary morphology 1, parameters I (A2, A3), capillary morphology 2, parameters II (B2, B3), and capillary morphology 3, parameters III (C2, C3). The amplification factor (maximal E_BBB_ per unit parenchyma electric field) were 367, 443, and 617, respectively for these three exemplary BBB capillary morphologies and parameters (A3; B3; C3). In addition, for each capillary morphology and variation, BBB resistivity (and so BBB space constant) was varied from a standard value (D1, E1; r_BBB_ = 1 × 10^5^ Ω.m) by a factor of 100 up (D2; E2; r_BBB_×100 = 1 × 10^7^ Ω.m) or down (D3; E3; r_BBB_/100 = 1 × 10^3^ Ω.m). For each FEM simulation, BBB polarization per unit brain parenchyma (BBB polarization length) and E_BBB_ per unit brain parenchyma (BBB Amplification factor) is summarized. Since E_BBB_ was not uniform across the vascular wall, we report “punctate” E*_BBB_ (at any point within the capillary wall) as well as the average E_BBB_ (V_BBB_ / d_th-BBB_). Finally, the analytically derived (see Methods) maximum BBB polarization length (λ_BBB_) and BBB Amplification factor (λ_BBB_/d_th-BBB_) is reported for each model.

**Figure 2:**
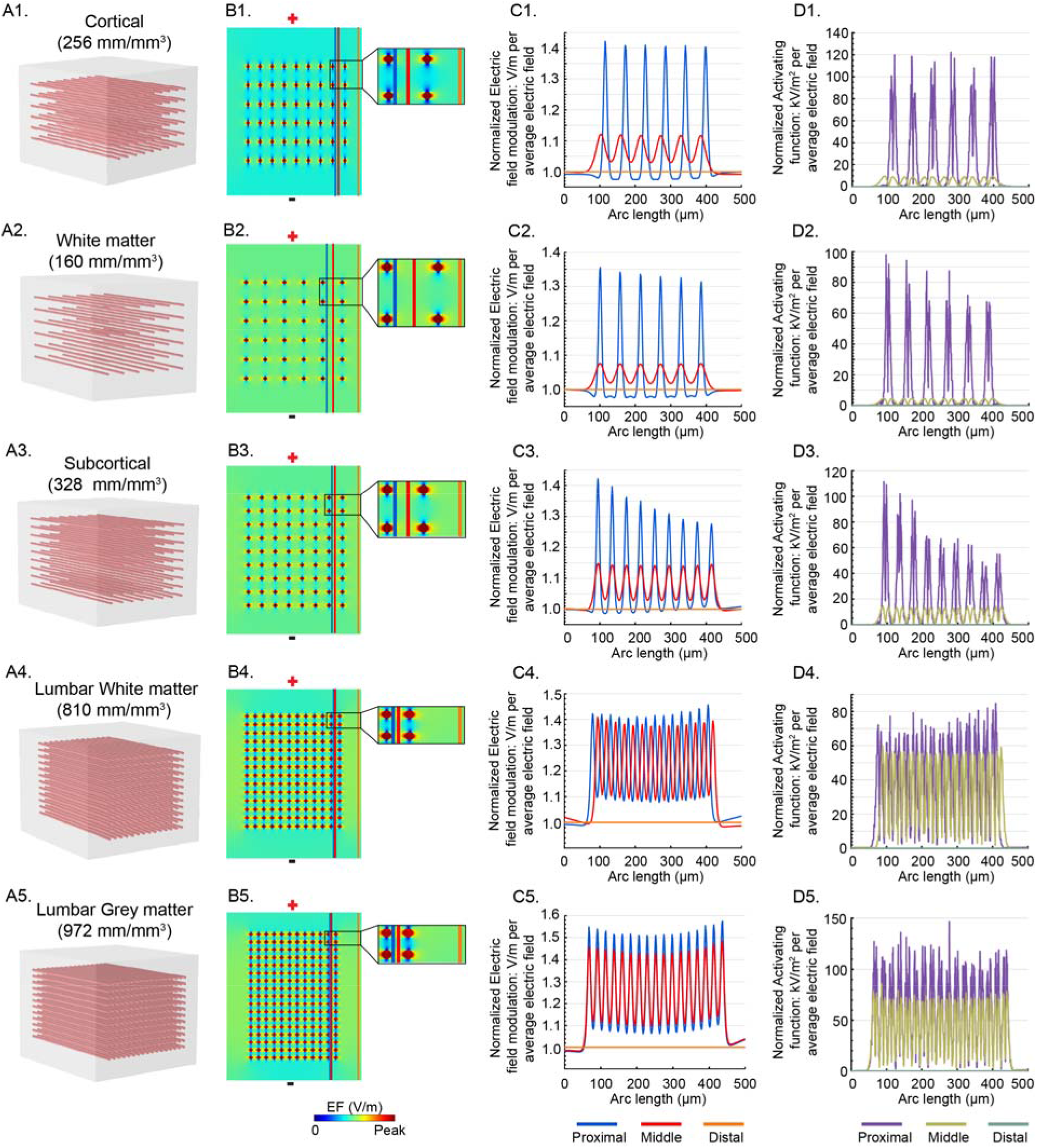
Impact of vascular ultrastructure on brain electric field. We consider vascular ultra-structure network for five brain regions (cortical, white-matter, subcortical, lumbar white-matter, and lumbar grey-matter). (A1, A2, A3, A4, A5) illustrates vascular network for these brain regions, noting the regional capillary length density (mm length per mm^3^ volume). (B1, B2, B3, B4, B5) Predicted electric field in a plane crossing the vascular bed, shows local distortion of electric field by the vasculature. Also illustrated is the straight trajectory for sampling of electric field and activating function: 1) Proximal trajectory (~ 5 μm away from nearest capillary; blue line), Middle trajectory (in between adjacent capillaries; red line), and Distal trajectory (region without capillary; orange line).(C1, C2, C3, C4, C5) Normalized electric field magnitude (per unit parenchyma electric field) along three trajectories. The degree of electric field modulation was higher for trajectories passing nearer capillaries and for denser capillary beds. (D1, D2, D3, D4, D5) Electric field gradient (Activating Function) magnitude (per unit parenchyma electric field) along three trajectories. Neuronal activation at the proximity of a vasculature was ~100 kV/m^2^ per unit average parenchyma electric field (Ē_BRAIN_). Activating functions were higher for trajectories passing nearer capillaries and for denser capillary beds.

Normal current density was applied to the one surface of the brain voxel while the opposite surface of the brain voxel was grounded, with the remaining external boundaries insulated. For computation, we used 0.08 A/m^2^ (corresponding to ~1 mA tDCS [38]), however all results were reported as normalized (i.e. per unit parenchyma electric field) by dividing results by the average (“bulk”) parenchyma electric field (Ē_BRAIN_). This is the same as the uniform electric field produced in a model with homogenous resistivity (i.e. only brain parenchyma). Laplace equation (∇·(σ∇V) = 0, where V is extracellular voltage and σ is electrical conductivity) was applied and solved as the field equation to determine the extracellular voltage distribution throughout the model. Three-dimensional (3D) extracellular voltage, electric field, and activating function were predicted in different capillary morphologies, and resulting BBB polarization length, BBB amplification factor, or neuronal polarization amplification by vascular ultrastructure were calculated.

### Models of BBB Electric Field Amplification (Principle 1): Numerical Solutions

For electric field amplification at the BBB, we simulated three variations of capillary morphology 1 namely I, I_1_, and I_2_, with fixed curvature width (1000 μm), and varied wall thickness (d_th-BBB_), lumen diameter (d_l_) and brain voxel volume. In variation I, the d_th-BBB_ was 10 μm, d_l_ was 100 μm, and brain voxel volume was 2.2 × 10^12^ μm^3^. In variation I_1_ and I_2_, the d_th-BBB_ was 1 μm and d_l_ was 10 μm, while the brain voxel volumes were 5.1 × 10^7^ μm^3^ and 1.1 × 10^8^ μm^3^, respectively. Unless otherwise mentioned, 2.2 × 10^12^ μm^3^ was used as a standard brain voxel volume for the remaining capillary morphology models. Capillary morphology 2 included two morphological variations namely II and II_2_. In both of these variations, the d_th-BBB_ was 10 μm and d_l_ was 100 μm, whereas the curvature widths were 1000 μm and 200 μm respectively for variation II and II_2_. Capillary morphology 3 included III and III_1_ morphological variations with variant terminal conditions. In variation III, one terminal of a semi-infinite straight tube was open, whereas both terminals were sealed in variation III_1_. The d_th-BBB_ was 10 μm and d_l_ was 100 μm for both III and III_1_ variations. The semi-circular loop of capillary morphology 1 and 2 or tapered end of capillary morphology 3 were oriented toward the energized surface the brain voxel. Capillary wall and lumen dimensions were based on cadaveric studies and imaging data [45–52].

Unless otherwise indicated, standard electrical resistivity (reciprocal of electrical conductivity) was assigned to each model domain as: capillary wall: 1 × 10^5^ Ω.m; capillary lumen: 1.42 Ω.m; and brain parenchyma: 3.62 Ω.m. In some simulations, capillary wall resistivity was increased or decreased 100-fold.

Capillary morphology 1 was positioned at the middle of the brain voxel in such a way that boundaries of capillary wall and lumen at the terminating ends of the orthogonal straight segments were sealed. Capillary wall and lumen boundaries at the terminating ends of the semi-infinite parallel segments of capillary morphology 2 were open (ground). Capillary morphology 3 was also positioned at the middle of the brain voxel, and the capillary lumen domain was enclosed by the capillary wall domain, with 1 μm spacing between them. Together they formed a semi-infinite membrane.

The numerical maxima for BBB polarization length (BBB polarization per unit parenchyma electric field) is given as:

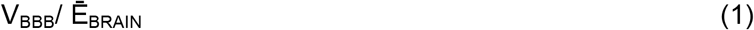

where V_BBB_ is a predicted BBB polarization (V) and (Ē_BRAIN_) is an average predicted parenchyma electric field (V/m). The numerically-computed average BBB electric field amplification (BBB electric field per unit parenchyma electric field) is expressed as:

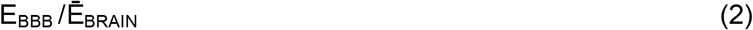

where E_BBB_ (V/m) is an average electric field across the capillary wall thickness, calculated as V_BBB_ per BBB thickness:

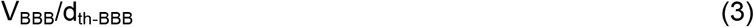

The punctate (maximal) BBB electric field amplification is expressed as:

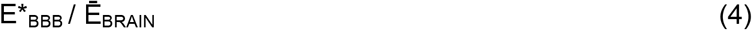

where E*_BBB_ (V/m) is the maximum predicted BBB electric field within the capillary wall, noting the electric field inside the capillary wall can change across the wall depth.

### Models of BBB Electric Field Amplification (Principle 1): Analytical Solutions

Analytical analysis of polarization of axon terminals in an electric field based on cable theory [43,44,53] shows the maximal polarization that can be experienced at a bent or terminating axon terminal as:

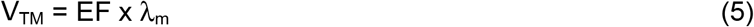

where V_TM_ is the change in axon terminal transmembrane potential, EF is the electric field around the terminal (V/m), and λ_m_ is the terminal space constant (m). λ_m_ is a function of only the axon membrane resistivity (r_m_: Ω.m) and axon intracellular resistivity (r_i_: Ω.m) as:

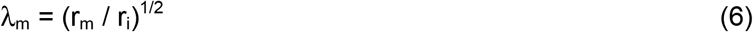

This maximal axon terminal polarization sensitivity may be secondarily amplified by “active” sub-threshold active channels at the terminal [54] and trigger a supra-threshold action potential. A maximal “passive” neuronal sensitivity of λ_m_ still applies, including to more complex neuronal structures [40,55].

Our analytical model for BBB polarization adapts this same cable theory where we model the capillary wall (BBB) as analogous to a continuous extracellular membrane and we model the capillary lumen (blood) as analogous to the continuous intracellular compartment. The analytically derived maximal BBB polarization is therefore expressed as:

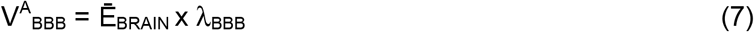

where V^A^_BBB_ is BBB polarization (V), Ē_BRAIN_ is an average parenchyma electric field (V/m), and λ_BBB_ is defined here as the BBB space constant (m). λ_BBB_ is a function of only the capillary wall (BBB) resistivity (r_BBB_: Ω.m) and capillary lumen (blood) resistivity (r_BLOOD_: Ω.m) as:

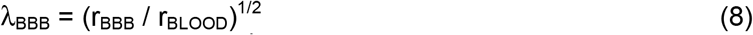

The analytical polarization length (V^A^_BBB_ per unit Ē_BRAIN_) is thus λ_BBB_. The maximal analytical BBB electric field is then expressed as:

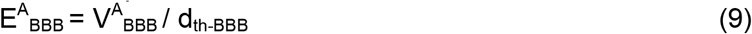

The analytical maximal amplification factor (E^A^_BBB_ per unit Ē_BRAIN_) is then estimated as:

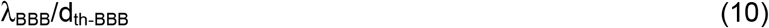

Blood vasculature structure and properties are not simply analogous to axons of neurons, so we use numerical FEM simulations of various exemplary capillary morphologies to test if our analytical solution predicts maximal BBB polarization and so also the maximal BBB electric field. While we designed the models such that the V_BBB_ and E_BBB_ were independent of brain voxel size, anomalous current patterns where blood vessel contacting model boundaries were not considered.

### Models of Neuron Polarization Amplification (Principle 2)

For neuron polarization amplification by vasculature, we modeled semi-infinite parallel solid capillaries, adjusting the length density (L_v_) of vasculature for varied brain regions (cortical grey-matter, white-matter, subcortical, lumbar white-matter, and lumbar grey-matter; Fig. 2) that are therapeutic targets (Table 1) for different modes of electrical stimulation (tDCS, TMS, ECT, DBS, and SCS). Solid capillaries were modeled with a uniform resistivity of 1 × 10^5^ Ω.m.

**Table 1:**
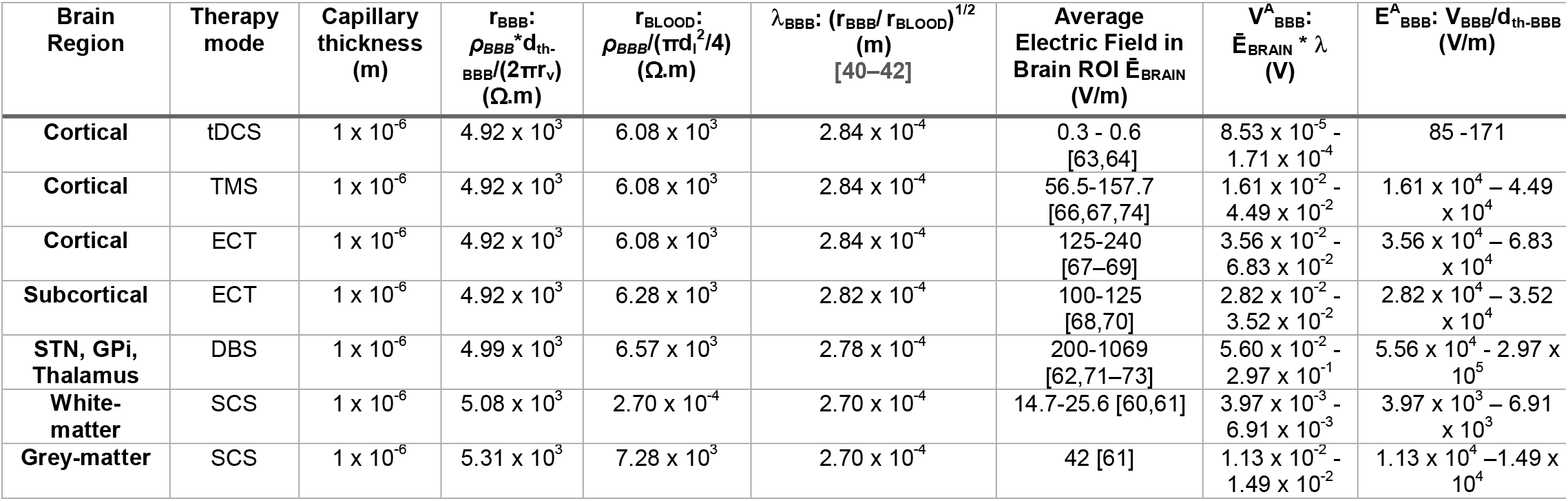
Predicted maximal V_BBB_ and E_BBB_ for various therapeutic modalities and brain targets. Region specific capillary anatomies and resistivities were used to calculate a representative BBB space constant (λ_BBB_) for each region. Based on our analytical derivation, maximum voltage across the BBB (V^A^_BBB_) and electric field across the BBB (E^A^_BBB_) is calculated.

Factors driving neuron polarization amplification by vasculature was quantified as normalized electric fields (per unit parenchyma electric field) and normalized activating functions (per unit parenchyma electric field) at three different brain voxel locations: proximal (~ 5 μm away from capillary), middle (in between two capillaries), and distal (no capillary zone) (Fig. 2B1-2B5).

## Results

### Theoretical Basis for Maximum Electric Field Amplification at the BBB (Principle 1)

To develop a theory quantifying BBB (vascular wall) polarization, resulting from current flow between the brain parenchyma and the blood during neuromodulation, we modeled stimulation across capillary segments of varied morphologies that are intended to capture maximum local polarization across a complex capillary network. We considered three prototypical capillary morphologies (Fig. 1 A1, B1, C1). Capillary morphology 1 was a semi-circular loop (fixed curvature width) with semi-infinite orthogonal straight segments, with variants of capillary size (I, I_1_, and I_2_). Capillary morphology 2 was a semi-circular loop (varied curvatures) with semi-infinite parallel straight segments with variants of loop curvature (II and II_1_). Capillary morphology 3 was a semi-infinite straight tube with two variants of terminal conditions (III, III_1_). FEM simulation predicted current flow though the brain voxel containing the capillary (Fig. 1 A2, B2, C2), and specifically current flow across the BBB (Fig. 1 A3, B3, C3). Models were designed so that maximum polarization was insensitive to the modeled tissue boundary size (see Methods).

For each morphology, the maximum voltage across the BBB (V_BBB_) and electric field across the BBB (E_BBB_) are reported as normalized to unit parenchyma electric field (E_BRAIN_). This allows reporting of BBB polarization length (V_BBB_ per unit E_BRAIN_; Fig. 1, row D) and the BBB amplification factor (E_BBB_ per unit E_BRAIN_; Fig. 1, row E). Thus, for any specific neuromodulation technology with a given average electric field in a brain target, the resulting BBB electric field is this average electric field times the region-specific amplification factor. Finally, for each capillary morphology, BBB resistivity was varied from a standard value (r_BBB_: Fig. 1D1, 1E1) up or down by a factor of 100 (r_BBB_×100: Fig. 1D2, 1E2; r_BBB_/100: Fig. 1D3, 1E3).

Note that the voltages (V_BBB_) and electric fields (E_BBB_) across the BBB segments varied for any capillary morphology; consistent with the objective of this section, we report local maxima for each stimulation. For example, peak E_BBB_ for the exemplary capillary morphologies I, II, and III (with standard r_BBB_) were, per unit Ē_BRAIN_: 367 V/m per V/m at capillary bend, 443 V/m per V/m at capillary bend, and 617 V/m per V/m at capillary terminal, respectively (Fig.1A3, 1B3, 1C3). We further predicted a varied electric field across the capillary wall thickness (i.e. the electric field changes across the BBB wall thickness). Unless otherwise stated, E_BBB_ is considered the average electric field across the capillary wall thickness for a given capillary segment, which is calculated using equation (3). In this section only, we also report the maximal “punctate” electric field across any point inside the capillary wall as E*_BBB_.

Based on cable theory (see Methods), we developed an analytical solution for maximum BBB polarization (V^A^_BBB_) which depends only on the space constant (λ_BBB_) of the vasculature (equation 7). When V^A^_BBB_ is expressed per unit Ē_BRAIN_, then the analytical maximum polarization length is simply λ_BBB_. The analytical solution for maximum BBB electric field (E^A^_BBB_) is then:

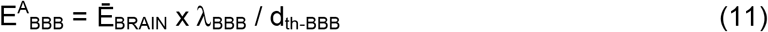

Thus, the analytical maximum electric field amplification factor is λ_BBB_ / d_th-BBB_.

For all the numerically (FEM) simulated capillary parameter, we also predicted (Fig. 1E, 1D) the corresponding analytical maximal BBB voltage (V^A^_BBB_) and electric field (E^A^_BBB_). λ_BBB_ depend on the square root of r_BBB_ (equation 8), as a result, V^A^_BBB_ and so E^A^_BBB,_ vary by 10x across 100x changes in r_BBB_. Note analytical predictions do not explicitly depend on capillary morphology (e.g. morphology 1, 2, or 3) but depend on BBB capillary wall and lumen properties. The I_1_ and I_2_ variations of capillary morphology 1 are thus the only models with different V^A^_BBB_ However, this difference is then absent for predicted E^A^_BBB_ because of additional dependence on d_th-BBB_ (equation 11).

In sum, across different variations of capillary morphologies and BBB capillary wall resistivities, we made two types of comparisons. First, for BBB polarization per unit parenchyma electric field, we compared numerical maxima (V_BBB_ per Ē_BRAIN_) with the analytical BBB polarization (V^A^_BBB_ per Ē_BRAIN_) based on λ_BBB_ (Fig. 1, row D). Second, for the BBB electric field amplification (BBB electric field per unit parenchyma electric field) we compared numerically-computed average (E_BBB_ per Ē_BRAIN_) and punctate (E*_BBB_ per Ē_BRAIN_) BBB electric field amplification with the analytical BBB electric field amplification (E^A^_BBB_ per Ē_BRAIN_) based on λ_BBB_ / d_th-BBB_ (Fig. 1, row E).

Across all simulated conditions, the numerically computed maximum polarization length (V_BBB_ per Ē_BRAIN_) was less than the analytical maxima (λ_BBB_). As a consequence, the numerically computed maximum average BBB electric field (E_BBB_ per Ē_BRAIN_) was also always less than the analytical maximum (λ_BBB_ / d_th-BBB_). In some models, the within-wall numerical maximum BBB electric field (E*_BBB_ per Ē_BRAIN_) exceed the analytical maximum, but never by more than by a factor of two. Provided our assumptions, the analytical solution for maximum BBB polarization (equation 7) and amplification factor (equation 10) can thus be considered reasonable approximations.

Finally, note that for Principle 1 analysis, an average (“bulk”) Ē_BRAIN_ was assumed, however distortion in electric field around the periphery of capillaries was already noted in these simulations and was central to the analysis of non-uniform E_BRAIN_ for Principle 2.

### Electric Fields Amplification at the BBB across Neuromodulation Interventions (Principle 1)

We considered five exemplary brain stimulation techniques (tDCS, TMS, ECT, DBS, and SCS) with associated brain targets (cortical, white-matter, subcortical, lumbar spinal white-matter, and lumbar spinal grey-matter). For each brain region, capillary anatomy (wall thickness: d_th-BBB_; capillary diameter: 2r_v_; lumen diameter: d_l_), and BBB membrane and blood resistivities (r_BBB_ and r_BLOOD_) were derived from prior literature [23–25,56–62]. These values were used to calculate a representative BBB space constant (λ_BBB_) for each brain region. Typical brain electric field produced by each stimulation modality were also derived from literature [63–73]. Finally, using the analytical method for predicting maximal BBB polarization length and BBB electric field amplification factor (Fig. 1), for each brain stimulation technique and associated brain region, the maximal BBB polarization (V_BBB_) and BBB electric field (E_BBB_) is predicted (Table 1).

The E^A^_BBB_ ranges from ~100 V/m for tDCS of cortex to ~100 kV/m for DBS. We note that variations in dose within each neuromodulation modality (e.g. electrode separation) and which brain region is considered (e.g. distance from electrode) causes E_BRAIN_ to vary. Moreover, E_BRAIN_ (and so E_BBB_) for any modality will vary linearly with applied current. Never-the-less, E^A^_BBB_ is consistently greater by over two orders of magnitude than Ē_BRAIN_. The temporal waveform of E_BBB_ would vary for each modality and programming as these setting effect E_BRAIN_. For example, E_BBB_ would be static for tDCS and would biophysically be pulse for other modalities. Our model assumes no temporal filtering (e.g. low pass) in the BBB amplification factor.

### Theoretical Basis for Neuron Polarization Amplification by Vascular Ultra-structure (Principle 2)

We developed a theory to predict distortion of current flow in the brain parenchyma by capillary ultrastructure and implications for maximum neuronal polarization. For cortical, white-matter, subcortical, lumbar spinal white-matter, and lumbar spinal grey-matter, we derived capillary bed length density (L_v_), surface density (S_v_), volumetric density (V_v_), numerical density (N_v_), and intercapillary distance (ICD) (Table 2). A representative vascular network of parallel solid capillaries was modeled for each brain region (Fig. 3, column A). The model was designed to be independent of brain voxel dimension and provide a conservative (uniform, no tortuosity) capillary distribution (see Methods).

**Table 2:** Vasculature network parameters of different brain region for various therapeutic mode of electrical stimulation.

| Brain Regions | Therapeutic mode | Length density ($L_v$: mm/mm<sup>3</sup>) | Surface density ($S_v$: mm<sup>2</sup>/mm<sup>3</sup>) | Volumetric density ($V_v$: mm<sup>3</sup>/mm<sup>3</sup>) | Numerical density ($N_v$: mm<sup>-3</sup>) | Intercapillary distance (ICD: $\mu\text{m}$)[75,76] |
| --- | --- | --- | --- | --- | --- | --- |
| Cortical | tDCS, TMS, ECT | 256 [77] | 7.9 | 0.02 | 492 | 45 |
| White-matter | TMS, ECT, DBS | 160 [78] | 4.9 | 0.01 | 307 | 57 |
| Subcortical | ECT, DBS | 328 [48,75,79] | 10.1 | 0.03 | 631 | 40 |
| Lumbar White-matter | SCS | 810 [80] | 24.9 | 0.06 | 1558 | 25 |
| Lumbar Grey-matter | SCS | 972 [80] | 29.9 | 0.07 | 1869 | 23 |

For each BBB geometry, the parenchyma electric field (E_BRAIN_) and electric field gradient (Activating Function) were calculated along three straight trajectories: Proximal (~5 μm away from a capillary at a nearest point), Middle (centered between adjacent capillaries, half the inter-capillary distance at a nearest point), and Distal (no capillary zone, ~100 μm from a capillary at a nearest point). E_BRAIN_ and Activating Function were reported (normalized to) per average parenchyma electric field (Ē_BRAIN_).

Electrical field in the brain parenchyma (E_BRAIN_) was distorted around brain vasculature, producing ~50% modulation of the average parenchyma electric field (Ē_BRAIN_) (Fig. 2, column B, column C). This change occurs within less than half of an inter-capillary distance, producing Activating Functions of ~100 kV/m^2^ per unit average parenchyma electric field (Ē_BRAIN_) (Fig. 2, column D). Both the depth of E_BRAIN_ modulation and spatial rate of change increased with capillary density.

### Neuronal Stimulation Driven by BBB Ultra-structure across Neuromodulation Interventions (Principle 2)

We considered five exemplary brain stimulation techniques (tDCS, TMS, ECT, DBS, and SCS) with associated brain targets (cortical, white-matter, subcortical, lumbar spinal white-matter, and lumbar spinal grey-matter). For each region, relevant capillary anatomy (Table 2) was used to calculate modulated E_BRAIN_ (the range of E_BRAIN_ changes) and Activating Function per unit average parenchyma electric field (Ē_BRAIN_). Next, we combined these constants with specific brain electric fields (Table 3). This analysis assumes negligible “macroscopic” change in E_BRAIN_ across brain voxel in the absence of vasculature (i.e. the electric field is uniform for a homogenous brain voxel) such that any local changes in E_BRAIN_ and non-zero Activating Function are introduced by the presence of vasculature. However, it is the macroscopic changes that are conventionally assumed to drive neuronal stimulation for many modalities. We thus, contrasted Activating Functions generated by conventional macroscopic tissue changes (values derived from literature; [35,61,67,68,70,73,81–85]) with the BBB ultra-structure generated Activating Function derived here. This comparison is subject to a range of assumptions (e.g. distance from electrodes) and simplifications (e.g. linear and homogenous capillary structure). Never-the-less, BBB ultra-structure driven changes may conservatively exceed those conventionally derived from macroscopic tissue changes (Table 3). Moreover, for some techniques, such as tDCS, the electric field is conventionally assumed uniform [35,38] (zero Activating Function), but our analysis instead suggest that it is non-uniform because of spatial modulation by BBB ultra-structure.

**Table 3:** Electric field modulation and Activating Function created by BBB ultra-structure for exemplary neuromodulation techniques and brain targets. E_BRAIN_ modulation and Activating functions are reported for the proximal neuronal trajectory.

| Brain Region | Therapy mode | Average Electric Field in Brain ROI $\bar{E}_{\text{BRAIN}}$ (V/m) | $E_{\text{BRAIN}}$ Modulation from Vascular Ultrastructure (V/m) | Neurovascular Activating Function (Vascular Ultrastructure) (V/m<sup>2</sup>) | Conventional Activating Function (Macroscopic) (V/m<sup>2</sup>) |
| --- | --- | --- | --- | --- | --- |
| Cortical | tDCS | 0.3 - 0.6 [63,64] | 0.15 – 0.3 | $1.10 \times 10^4 - 2.21 \times 10^4$ | 0 ([35,81]) |
| Cortical | TMS | 56.5-157.7 [66,67,74] | 28.3 – 78.9 | $6.23 \times 10^5 - 5.80 \times 10^6$ | 0 ([67,70]) |
| Cortical | ECT | 125-240 [67–69] | 62.5 - 120 | $4.60 \times 10^6 - 8.82 \times 10^6$ | 0 ([68,85]) |
| Subcortical | ECT | 100-125 [68,70] | 50 – 62.5 | $3.35 \times 10^6 - 4.19 \times 10^6$ | 0 ([68,85]) |
| **STN, GPI, Thalamus** | DBS | 200-1069 [62,71–73] | 100 – 534.5 | $6.70 \times 10^6 - 3.58 \times 10^7$ | $2.2 \times 10^5$ ([73,82]) |
| **White-matter** | SCS | 14.7-25.6 [60,61] | 7.4 – 12.8 | $3.74 \times 10^5 - 6.52 \times 10^5$ | $4.70 \times 10^4$ ([61,83]) |
| **Grey-matter** | SCS | 42 [61] | 21 | $1.85 \times 10^6$ | $8.14 \times 10^3$ ([61,83]) |

## Discussion

The study of which neural elements are activated by neuromodulation is exhaustive and includes verification in isolated systems without vasculature [86–88]. The first principle of neurovascular-modulation, that primary stimulation of BBB function leads to secondary changes in neuronal activity, is complimentary to these conventional theories of direct neural stimulation. We predict the maximal electric field across the BBB (E_BBB_) are over two orders of magnitude above brain parenchyma (E_BRAIN_), with a maximum amplification factor (λ_BBB_/d_th-BBB_) adapted from the cable theory. Electric field across the BBB modulate water and solute transport [20–22] which in turn regulate neuronal metabolic capacity and interstitial clearance. Brain imaging techniques that depend on hemodynamic changes are a bedrock of systems neuroscience (e.g. fMRI, fNIRS) - we suggest that in the specific case of neuromodulation they can be interpreted as suggestive of direct vascular modulation (first principle) rather than secondary neurovascular coupling.

Brain hemodynamics (neurovascular coupling) and BBB transport are disrupted in brain disease, including Alzheimer’s Disease and Parkinson’s [89–91] and following brain injury [7,92,93]. Indeed, BBB dysfunction may be a link across these disorders [94,95]. Notably, while Alzheimer’s disease is traditionally considered a disease of neurofibrillary tangles and amyloid plaques, structural and functional changes in the microvessels may contribute directly to the pathogenesis of the disease [96–100], specifically disruption of brain clearance systems dependent on (water) transport across the BBB [29,101,102]. For a wide range of brain disorders, there is interest in interventions modulating brain hemodynamics and clearance system; neuromodulation may have powerful and unique actions (Principle 1).

When neuromodulation drives intense neuronal activity or relies on neuroplasticity, then neuromodulation is governed by brain metabolism, and so by neurovascular dynamics. The direct stimulation of the BBB by neuromodulation (Principle 1) may thus also play a role in modulating metabolically active states created by direct neuronal stimulation mechanisms. To the extent hemodynamic based functional imaging of neuromodulation does not reflect direct BBB stimulation (Principle 1) but rather conventional neurovascular coupling, it still reinforces the role of the BBB in governing neuronal responses.

The second principle of neurovascular-modulation address direct neural stimulation but with efficacy that is governed by current flow distortion around vascular ultra-structure. We develop a theory relating capillary density to local fluctuations in E_BBB_. Stimulation of neurons is traditionally modeled as reflecting two cases: 1) changes in E_BRAIN_ along the neural structure (Activating Function; [44,103,104] or 2) polarization by locally uniform E_BRAIN_ [40,55,105]. In the first case, E_BRAIN_ gradients are conventionally assumed to reflect macroscopic variation in both tissue resistivity and decay with distance from electrodes. However, by Principle 2, local E_BRAIN_ gradients produced by BBB ultra-structure may overwhelm those changes driven by traditional macroscopic models (Table 3). In the second case, Principe 2 suggest locally uniform brain electric fields may in fact not exist. In both cases, that stimulation dose and macro-tissue properties still govern the “incident” E_BRAIN_ arriving at each brain target (modeled here as the average parenchyma electric field (Ē_BRAIN_)), which is then modulated by regional BBB ultra-structure. In this sense, the quasi-uniform assumption remains valid [38,39,106].

These neurovascular-modulation principles are unrelated to BBB injury by electrical stimulation which depends on electrochemical products [107,108]. Activation of neurogenic regulation of cardiac function [109–111] or brain clearance [112] including electrical stimulation of perivascular innervation [113] is distinct from the direct BBB polarization of Principle 2. Electrical stimulation of glia [114–116] and subsequent astrocyte regulation of the BBB [117] are also parallel but distinct pathways.

The capillary bed of the brain is comprised of a dense network of intercommunicating vessels formed by specialized endothelial cells. Endothelial cells and pericytes are encased by basal lamina (~30 – 40 nm thick) containing collagen type IV, heparin sulfate proteoglycans, laminin, fibronectin, and other extracellular matrix proteins [118]. The basal lamina of the brain endothelium is continuous with astrocytic end-feet that ensheath the cerebral capillaries [119,120]. None of these details were modeled here and point to still more intricate mechanisms of neurovascular-modulation.

## Acknowledgments

Source(s) of financial support: This study was partially funded by grants to MB from NIH (NIMH 1R01MH111896, NINDS 1R01NS101362, NCI U54CA137788/ U54CA132378, R03 NS054783, 1R01NS112996-01A1) New York State Department of Health (NYS DOH, DOH01-C31291GG), and cycle 50 PSC-CUNY.

